# Protective effect of P2Y receptors antagonism on stress-induced retinal degeneration

**DOI:** 10.1101/2024.05.28.596247

**Authors:** Yi Bao, Kyle Bond, Pauline Sarraf, Robert Esterberg, Megan Serpa, Michael Twarog, YongYao Xu, Heather MacLeod, Qian Huang, Magali Saint-Geniez

## Abstract

The death of retinal pigment epithelial (RPE) cells and photoreceptors (PR) is a hallmark of the progression of several degenerative ocular disorders. The precise molecular driver(s) behind RPE and PR cell death, however, remains unknown. Recent studies have suggested the involvement of ATP and purinergic signaling in the progression of age-related macular degeneration (AMD) and retinal degeneration. We have discovered that RPE cells release ATP when subjected to stress, which in turn exacerbates stress-related signaling via purinergic receptors that ultimately results in degeneration. Our findings demonstrate that blocking P2Y purinergic receptors using suramin can effectively prevent toxin-induced RPE cell death and dysfunction *in vitro*. Furthermore, we show efficacy of suramin in preventing photoreceptor degeneration *in vivo* using the RHO-P23H zebrafish model. This study reinforces the involvement of ATP and purinergic signaling in maintaining retinal health, and highlights the potential of purinergic receptor antagonism as a therapeutic strategy for retinal degeneration.

## Introduction

Changes in health of the retinal pigment epithelium (RPE) and photoreceptors (PR) is hallmark in the progression of several degenerative eye diseases, such as age-related macular degeneration (AMD), Stargardt’s disease, and retinitis pigmentosa (RP) (1-4). The PR, consisting of rods and cones, are the primary drivers of visual function in the retina. Rod and cone cells contain the light sensing receptors rhodopsin (RHO) and opsin, respectively, which initiate light signal transduction. Within the PR outer segments (OS), the chain of biochemical reactions that contribute to the visual cycle are energetically demanding and generate many toxic metabolites. OS require clearing and regeneration to maintain homeostasis. Disruption of this homeostasis, or lack of RHO and opsin, is associated with genetic degenerative diseases such as RP. Continuous stress to the PR can lead to cell death and unrecoverable vision loss. Also responsible for the maintenance of visual cycle homeostasis are the RPE, lying at the interface between PRs and the choroid, which are essential for phagocytosing shed OS and recycling visual metabolites to replenish the neural retina (5-7). The RPE are central to the visual cycle by supporting the health of PR via nutrient exchange and clearance of toxic metabolites through phagocytosis of the OS. Genetic risks and environmental factors can promote RPE dysfunction and atrophy, ultimately leading to retinal degeneration (8-11). Disruption of any of the myriad homeostatic functions of RPE can result in retinal degeneration; however, the specific nature of their involvement in ocular disease continue to be elucidated (11,12).

Beyond energy currency, adenosine triphosphate (ATP) is a crucial signal molecule involved in many physiological processes. Within the retina, ATP is critical to the photoransduction cascade responsible for vision. It also serves as a key mediator of cell signaling in purinergic signaling that involves purine nucleotides and their breakdown products within extracellular space. Extracellular ATP can regulate cell functions by stimulating a large family of purinergic receptors that are widely expressed and found on all mammalian cell types. Purinergic receptors include three different families: the P2X, P2Y, and P1 receptors. Mammalian cells can express various combinations of the seven P2X receptors that act as ATP-gated ion channels, the eight known P2Y receptor subtypes that belong to the G protein-coupled receptor (GPCR) superfamily, and the four different adenosine (P1) receptors that are GPCRs and can couple to Gαi (A1 and A3) or Gαs subunits (A2A and A2B). The diversity of the purinergic receptor subtype profiles that can be present in mammalian cells allows purinergic signaling mechanisms to promote or inhibit cell responses through the different downstream signaling events they can trigger (13-15).

In recent years, the role of ATP and purinergic signaling in the retina has garnered significant attention due to its implications in retinal function, degeneration, and disease. In the retina, ATP acts as a potent neurotransmitter and modulator, participating in purinergic signaling pathways that regulate various aspects of retinal function. Components of the visual cycle, most notably the opsin photopigments, initiate light-sensitive signaling cascades through their GPCR activity rely upon proper ATP balance to function properly. ATP levels are disrupted in human retinal disease and in preclinical degeneration models, raising the possibility that they contribute in some way to the process. For example, significantly higher ATP levels have been reported in the vitreous humor of AMD patients compared to non-AMD donors (16). Furthermore, Mitchell et al determined ATP can be released from RPE cells via gap junction hemichannels to the membranes facing the neural retina, and that this release of ATP is regulated by the intracellular calcium concentration (17,18). It was also reported that, once released, extracellular ATP may cause further RPE cell death and retinal degeneration via P2X7 and inflammasome activation (19).

In this study, we found that ATP was released from RPE cells under stress or death. Additionally, released ATP can intensify RPE damage induced by other stressors. Finally, we show that preventing purinergic signaling using a pan-purinergic receptor antagonist, suramin, can alleviate stimuli-induced RPE and PR degeneration. Together, these data suggest that an abundance of extracellular ATP negatively influence disease trajectory, and therapies aimed at returning extracellular ATP to homeostatic levels may be beneficial during degeneration.

## Methods

### Cell Culture

The human RPE cell line, ARPE-19, was purchased from ATCC (Manassas, VA; CRL-2302). ARPE-19 were cultured in DMEM/F12 (Gibco, Carlsbad, CA) supplemented with 10% heat-inactivated fetal calf serum (FBS) (Sigma-Aldrich, St. Louis, MO) and 1% penicillin/streptomycin (Gibco) at 37 °C and 5% CO_2_. Cells were seeded at high density and maintained for at least 3 weeks to form mature monolayers, unless otherwise specified.

### Reagents

All-trans-retinal (atRAL) was purchased from Toronto Research Chemicals (TRC, Canada), and stored or manipulated with limited light exposure. 4-Hydroxynonenal (4-HNE) was purchased from Millipore Sigma (Burlington, MA). ATP, apyrase and reactive blue 2 (RB-2) purchased from Sigma. Suramin, PPADS, MRS2500, AR-C118925xx and CBX purchased from Tocris Bioscience (Minneapolis, MN).

### Measuring and visualizing ATP

The supernatants were collected from cells treated with indicated stimulations, and the amount of ATP released was determined using a commercially available luciferase-based ATP Determination Kit (Invitrogen). Dose response curve and relevant controls were applied to confirm the kit works well for measuring ATP levels (Supplementary Figure 1). 2-2Zn(II) was applied for visualizing membrane bound ATP (20). Cells were incubated with 1 µM 2-2Zn(II) for 5 min, then images were acquired using Zeiss confocal microscope. Images were analyzed using Fiji.

### Measuring ARPE-19 cell death and stress

To induce cell death in ARPE-19 cells, various methods were employed. One approach involved exposing the cells to specific toxins such as atRAL or 4-HNE to induce global cell death. Another method involved utilizing Zeiss LSM 880 confocal 405 nm laser with 100% power to bleach a focal region of cells for a duration of 30 minutes, with intervals of 30 seconds. This laser treatment was designed to inflict targeted damage and induce localized cell death. Images were obtained using a 20x magnification on an LSM 880 microscope, employing z or t stacks to capture multiple focal planes or time points, respectively.

Propidium iodide (PI), red-fluorescent nuclear and chromosome counterstain for dead cell detection, was purchased from Thermo Fisher Scientific (Waltham, MA). Cell death was measured following manufacturer’s protocol. Fluorescent signals were visualized and recorded as time-lapse images using IncuCyte (Sartorius).

### Cell morphology evaluated by ZO-1 staining

Cells were fixed with 4% paraformaldehyde for 15 min, incubated in 3% bovine serum albumin, 0.1% Tween-20, and 0.1% Triton-X-100 for 1 h, then stained with ZO-1 antibody (Invitrogen) at 4°C for 18 h. Samples were washed and incubated with Alexa Fluor 647-conjugated secondary antibody (Invitrogen) and DAPI (Invitrogen) for an additional 30 min at room temperature. Images were acquired using 20x magnification on a Molecular Devices ImageXpress Micro imager.

### Barrier function evaluated by impedance

Impedance experiments were carried out using the xCelligence Real Time Cell Analysis (RTCA) platform (Acea Biosciences, California). ARPE-19 cells were seeded in each well (E-plate 96, ACEA Biosciences) and cultured until confluent. Cells were incubated with suramin 1 h prior to atRAL or 4-HNE challenge and impedance measured for at least 24 h after challenge. The impedance value of each well was automatically monitored by the xCELLigence system and expressed as a CI (cell index) value. Data for cell impedance were normalized to the value at the time of ligand addition. Normalized CI is calculated using the software provided by the vendor, end time point reading was used for calculation of “% of increase”, which is equal to (CI of compound with ligand-CI of media)/(CI of ligand alone-CI of media)*100%.

### Zebrafish model generation

All procedures employing animals have been reviewed and approved by the Institutional Animal Care and Use Committee at Novartis Biomedical Research. Zebrafish (Danio rerio) breeding and care were performed according to standard methods (21). The AB strain was maintained on a 14/10 light-dark cycle. Randomly selected larval and juvenile fish of both sexes were used for experiments.

Transgenic animals were generated using the Tol2 Gateway system (22). A 5’ entry vector was generated containing 5.5 kb fragment of the Xenopus rhodopsin promoter to drive rod-specific expression as previously described (23). A middle entry construct was generated using human RHODOPSIN CDS lacking a stop codon, with a c.68 C>A variant generated to facilitate Pro>His amino acid substitution. A 3’ entry vector was generated containing a P2A self-cleavage site (24) and RFP. Entry constructs were synthesized through an outside vendor containing appropriate (ie, att) sites to facilitate cloning into the pDestTol2 destination vector according to manufacturer instructions and as previously described (22).

Tol2 transposase mRNA made by in vitro transcription of the pCS2+ plasmid using the mMessage mMachine kit (Life Technologies, Austin, TX, USA), together with transgene plasmid DNA were co-injected into 1-cell stage embryos. The resulting fish were grown to adulthood, outcrossed to transgenic AB zebrafish line containing a GFP transgene containing a CCIIL in-frame fusion at the C terminus to enable labeling of rod outer segments (25). Pools of embryos were screened for transgene transmission through PCR using M13 Forward or Reverse primers unique to the construct and/or the presence of the RFP transgene. Offspring exhibiting strong transgene expression were expanded through outcrossing to the AB strain, and used for studies described here. Transgenic fish were genotyped by PCR of tail cut DNA as described elsewhere (26).

### *In vivo* dosing and analysis

For dosing experiments, animals at 3 days post-fertilization (dpf) were dosed via immersion using suramin dissolved in 0.01% DMSO and E3 embryo medium at final concentration of 125 µM, at which concentrations did not elicit adverse effects (data not shown). Media containing compound were refreshed daily. Following treatment, embryos were euthanized at 7 dpf using Tricaine, embedded in OCT mounting media to enable transverse sectioning through the retina, and sectioned at 10 µm thickness using standard methods (27). Only transverse sections through the optic nerve were included for photoreceptor quantification. Anti-GFP antibodies (Abcam; ab13970) at a dilution of 1:500 were used to amplify transgenic markers according to standard immunofluorescence methods (27). Rods were quantified manually as a function of GFP+ cells.

### Data analysis

For all experiments, at least three independent experiments with technical triplicates were performed, and values were presented as mean±S.D.. Non-parametric statistic methods were applied. For two groups were compared using Mann-Whitney test, and for multiple comparisons were performed using Kruskal-Wallis test by GraphPad Prism. Differences were considered significant at * *p*<0.05, ** *p*<0.01 and *** *p*<0.001.

## Results

### ATP is released from RPE under stress and cell death

The RPE have specialized functions that, when impaired, can contribute to progressive RPE degeneration. Deficiency in key enzymes and/or exposure to excessive light leads to disrupted clearance and accumulation of toxic intermediate metabolites generated by the visual cycle, such as all-trans-retinal (atRAL) (1,28). atRAL was found to be cytotoxic in both rat and human RPE cells (29-31). To understand the role of purinergic signaling in atRAL mediated cell death, we measured ATP secretion from ARPE-19 cells, a spontaneously arising human RPE cell line, after treatment with atRAL using a luciferase-based ATP Determination Kit. A low level of ATP is released from ARPE-19 under normal conditions (∼1-2 nM Figure 1A). Vehicle treatment caused about 10-20 nM ATP release from ARPE-19 cells, which is consistent with the amount of ATP release due to mechanical stimulation (Figure 1A, Supplementary Figure 1B). However, within 30 min of stimulation with 18 µM atRAL, RPE cells began to secrete higher levels of ATP (∼ 40 nM), indicating that metabolic stress may induce ATP release preceding cell death (Figure 1A).

**Figure 1.**
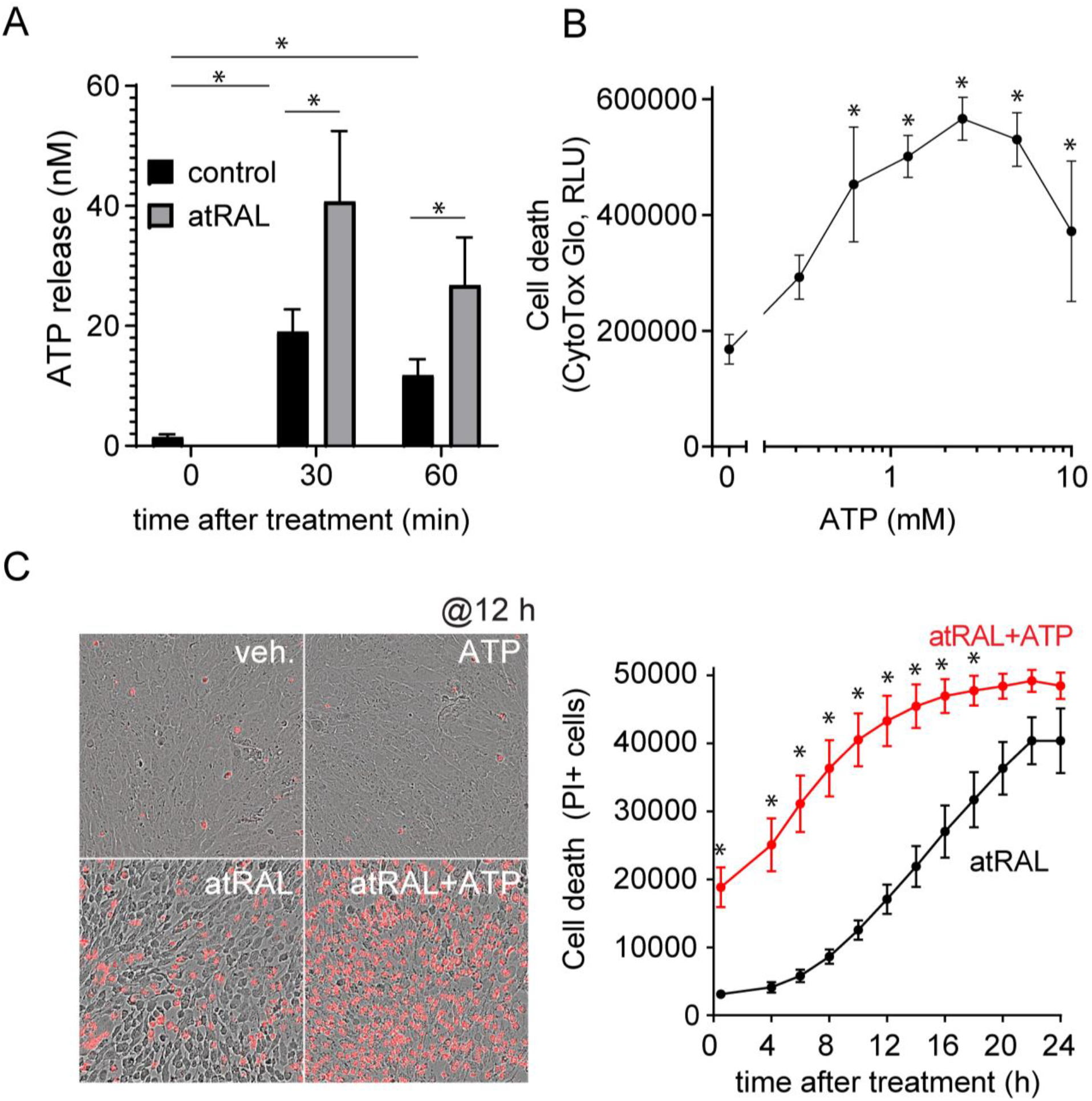
Stress-induced ATP release causes cell death and enhances atRAL-induced cell degeneration in ARPE-19. (A) atRAL induces ATP release from ARPE-19. ARPE-19 cells were trated with atRAL (18 µM) at indicated time points, and the amount of ATP release was evaluated in the supernatant using an ATP measurement kit. (B) ATP causes ARPE-19 stress and cell death. Indicated concentrations of ATP were applied on ARPE-19 cells for 24 h, and activity of protease released from stressed or dead cells in the supernatant was evaluated using CytoTox-Glo. (C-D) ATP enhances atRAL-induced ARPE-19 cell death. ARPE-19 cells were treated with atRAL (18 µM), or combine with ATP (100 µM), and cell death was measured by PI staining and imaged for 24 h using the IncuCyte. Represented images at 12 h after stimulation was shown in left panel of Figure 1C. PI+ cells were counted and evaluated by IncuCyte software (D). Data presented as mean±S.D., * *p*<0.05 compared to control.

While ATP may be released in response to stress-stimulation, it may also be released during cell death. To delineate the role of ATP during progressive cell death *in vitro*, we developed a system to measure ATP secretion from ARPE-19 cells after cell damage. Using a laser, we were able to selectively induce death in a discreet region of a monolayer of ARPE-19, similar to the localized atrophic regions characteristic of late-stage AMD (32,33). The fluorescently conjugated ATP-specific binding metal ion Green-2-2Zn(II) (2Zn) was utilized to visualize ATP (20). A basal level of ATP was observed on the ARPE-19 cell surface, and the addition of extracellular ATP or apyrase, which catalyzes hydrolysis of ATP, confirmed the selectivity of the dye (Supplementary Figure 2A-B, Supplementary Movie 1). Cell death was rapidly induced within the subjugated region following laser exposure, as indicated through propidium iodide staining. Interestingly, within 30 min, secondary cell death was observed in cells surrounding the initial laser exposed site. Additionally, significant levels of ATP were observed in both laser-induced death and secondary cell death within the same time window (Supplementary Figure 2C-D, Supplementary Movie 2). We hypothesized that this secondary cell death was due to the release of ATP from the initial attack, causing an increase in the “lesion” growth. We conclude that ATP secretion by RPE can happen at moderate levels in response to acute stress, or at high levels following death.

### ATP intensifies toxin-induced RPE cell death

Purinergic signaling within the subretinal space is important; activation of P1 and P2 receptors on RPE cells controls phagocytosis and fluid absorption, while activation on the photoreceptors can induce apoptosis (16). Since ATP is known to be an initiator of cell death, but also released because of it, we evaluated whether ATP could compound atRAL-induced cell death. Similar to previous reports (19), we observed a minimal concentration of 10 mM ATP could directly induce ARPE-19 cell death, as measured through PI staining (Supplementary Figure 3). Using CytoTox-Glo kit to measure protease release from cells under stress and cell death, we observed high levels of the protease release following 100 µM ∼ 2 mM of ATP exposure as well (Figure 1B), indicating cell under stress and ultimatly cell death. In addition, when combined with atRAL we observed a significant increase in cell death rate with lower dose of ATP (100 µM) compared to atRAL alone (Figure 1C-D), indicating that purinergic signaling sensitizes RPE cells to further damage or death. We conclude that ATP is not only released by cells under stress, but exacerbates it.

### Suramin prevents RPE cell death via inhibition of P2Y receptors

Purinergic receptors are essential to initiate downstream signaling in response to extracellular ATP.To investigate if purinergic targeting could serves as approach for intervention, we utilized suramin, a well-documented pan-P2 receptor antagonist, to inhibit purinergic signaling. Suramin dose-dependently reduced atRAL-induced ARPE-19 cell death, and fully prevented death at high doses (>100 µM) (Figure 2A-B). In addition, we also evaluated if modification of the treatment window influenced the protective effects, i.e. if intervention before or after toxin exposure could recover cell loss. Interesting, no matter the protection model (pretreatment of suramin) or therapeutic model (post-treatment of suramin after toxin induction), suramin effectively prevented atRAL induced ARPE-19 cell death (Figure 2C). Previously, reports suggested that different RPE specific stressors may induce different forms of cell death. While atRAL induces apoptosis, the oxidative byproduct 4-hydroxynonenal (4-HNE) induces a necrotic type of cell death, both of which may be involved in RPE degeneration and AMD progression (12,34,35). Surprisingly, despite the distinct differences in modality, suramin could also prevent 4-HNE induced RPE cell death (Figure 2D).

**Figure 2.**
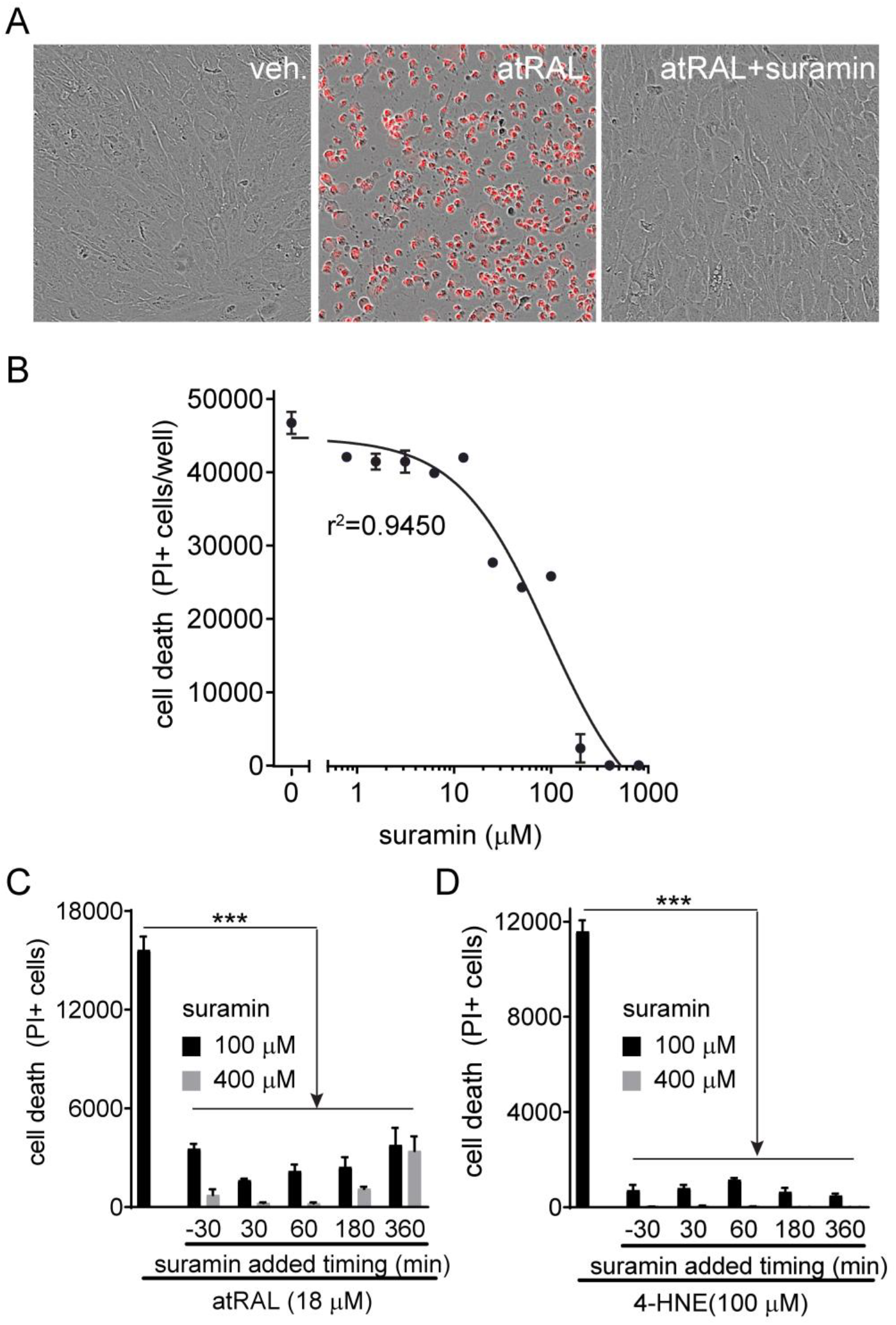
Blocking purinergic signaling using suramin prevents atRAL- or 4-HNE-induced ARPE-19 cell death. (A-B) Suramin prevents atRAL-induced ARPE-19 cell death. ARPE-19 cells were pretreated with the indicated doses of suramin or vehicle, then exposed to atRAL (18 µM). Cell death was measured using PI staining. Represented images are shown (A) and the dose response curve was calculated (B). (C-D) Suramin prevents atRAL-(C) or 4-HNE- (D) induced ARPE-19 cell death in both a preventative (pretreatment of suramin before stimuli) and therapeutic (post treatment of suramin after stimuli) treatment model. Data presented as mean±S.D., *** *p*<0.001 compared to relevant control.

To better understand how purinergic signaling contributes to cell death, we investigated several known inhibitors of purinergic signaling. This included the connexin/pannexin channel blocker carbenoxolone (CBX) to inhibit ATP release, and different purinergic receptor inhibitors to better distinguish which receptor may involve in this process in RPE. At high concentrations (100 µM), CBX showed some protection against atRAL mediated cell death, however, equal concentrations of suramin almost entirely prevented cell death (Figure 3A-B). P2 receptors can be divided into two subtypes; the ionotropic P2X and the metabotropic P2Y receptors. Suramin has been reported to block both. To delineate the importance of individual P2 receptors, atRAL treated cells were co-treated cells with suramin, the pan-P2X receptor antagonist PPADS, or the pan-P2Y antagonist RB-2. In a dose dependent manner, cells were protected from atRAL mediated cell death when treated with suramin as well as RB-2 (Figure 3C-D). However, no protective effects were observed with PPADS (Figure 3C-D), indicating that the protective role of suramin may be through blockade of P2Y receptor activation. Interestingly, inhibiting specifically P2RY1 or P2RY2 using inhibitors MRS2500 and AR-C118925xx, respectively, did not show the same protective effect as suramin did (Figure 3E-F), suggesting P2Y receptors may perform redundant signaling roles, and a pan-P2Y inhibitor may be needed to inhibit all P2Y downstream signaling to reach the full protective effects.

**Figure 3.**
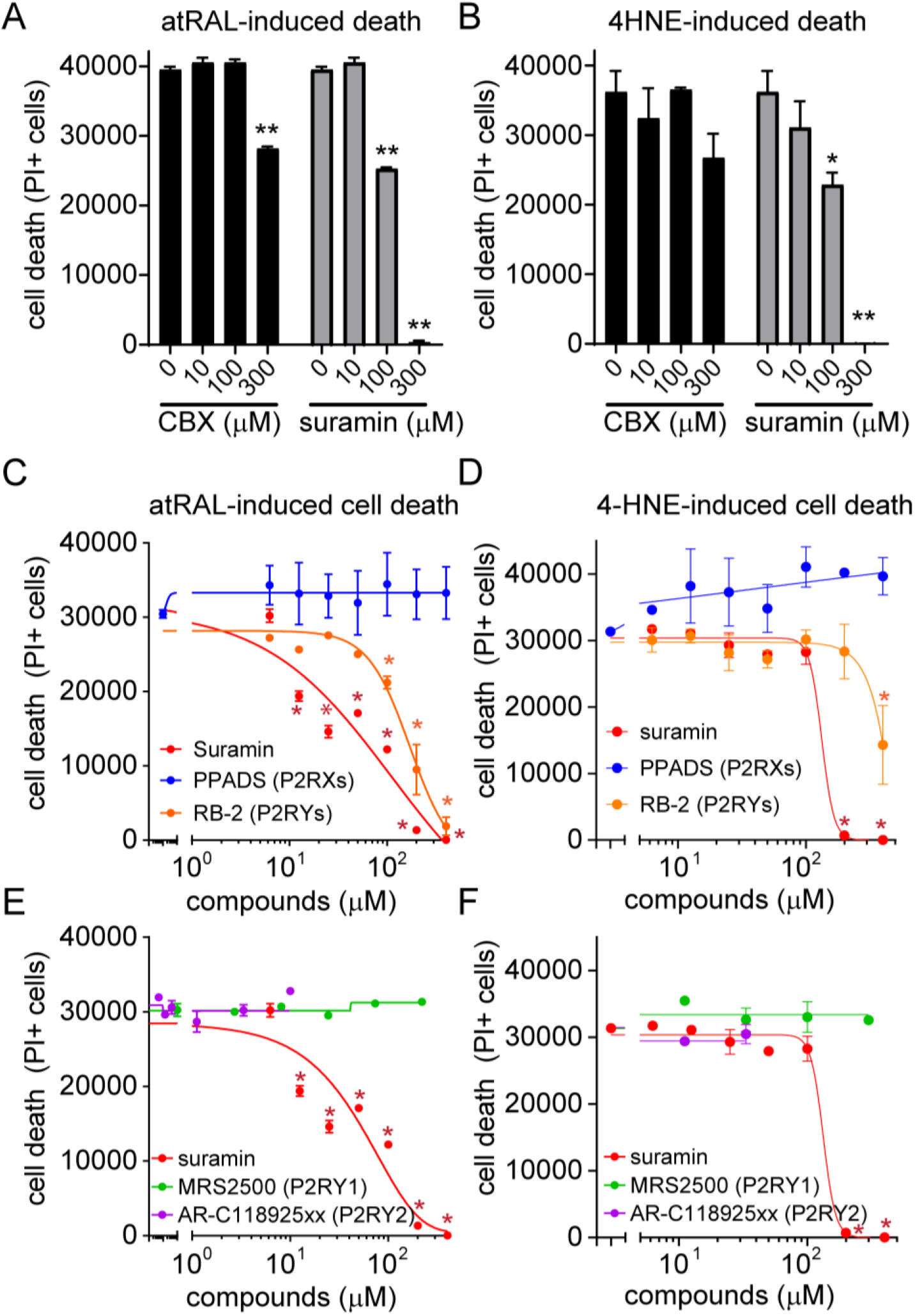
Suramin prevents toxin-induced ARPE-19 cell death via P2RY receptor inhibition. (A-B) Inhibition of pannexin-1 channels in ARPE-19 by CBX at different doses partly prevents atRAL- (A) or 4-HNE- (B) induced cell death. Cell death was measured by PI staining. (C-D) Inhibition P2RY but not P2RX prevents toxin-induced cell death. Pan-P2RY and pan-P2RX inhibition in ARPE-19 using varying doses of PPADs, RB-2, and suramin with co-exposure to atRAL (C) or 4-HNE (D). Cell death was measured by PI staining. (E-F) Blockade of specific P2RY receptors is not sufficient to prevent toxin-induced cell death. Specific inhibition of P2RY1 and P2RY2 in ARPE-19 using varying doses of MRS2500, AR-C118925xx, and suramin with co-exposure to atRAL (E) or 4-HNE (F). Cell death was measured by PI staining. Data presented as mean±S.D., * *p*<0.05, ** *p*<0.01 compared to relevant control.

### Suramin protects cells from toxin induced RPE stress and dysfunction

Continuing our exploration on the protective effects of suramin on RPE cells, we investigated whether suramin could restore RPE functionality following stressor-induced injury. Healthy RPE cells form an impermeable monolayer of cells organized by tight junctions that, when disrupted, serves as a hallmark of dysregulation. By measuring impedance, we can evaluate barrier function loss due to stress induction. We show that both atRAL and 4-HNE reduce impedance levels in ARPE-19 cells. Excitingly, treatment with suramin prevented toxin-induced impedance loss under both conditions (Figure 4A), suggesting functional protection. Further investigation of tight junctions through staining for the scaffolding protein, ZO-1, confirmed that while atRAL and 4-HNE caused disruption of the monolayer and loss of compact tight junctions between cells, tight junctions were maintained in patches of cells exposed to suramin (Figure 4B-C). From this we conclude that suramin has highly advantageous effects on RPE health and functionality in response to stress induction.

**Figure 4.**
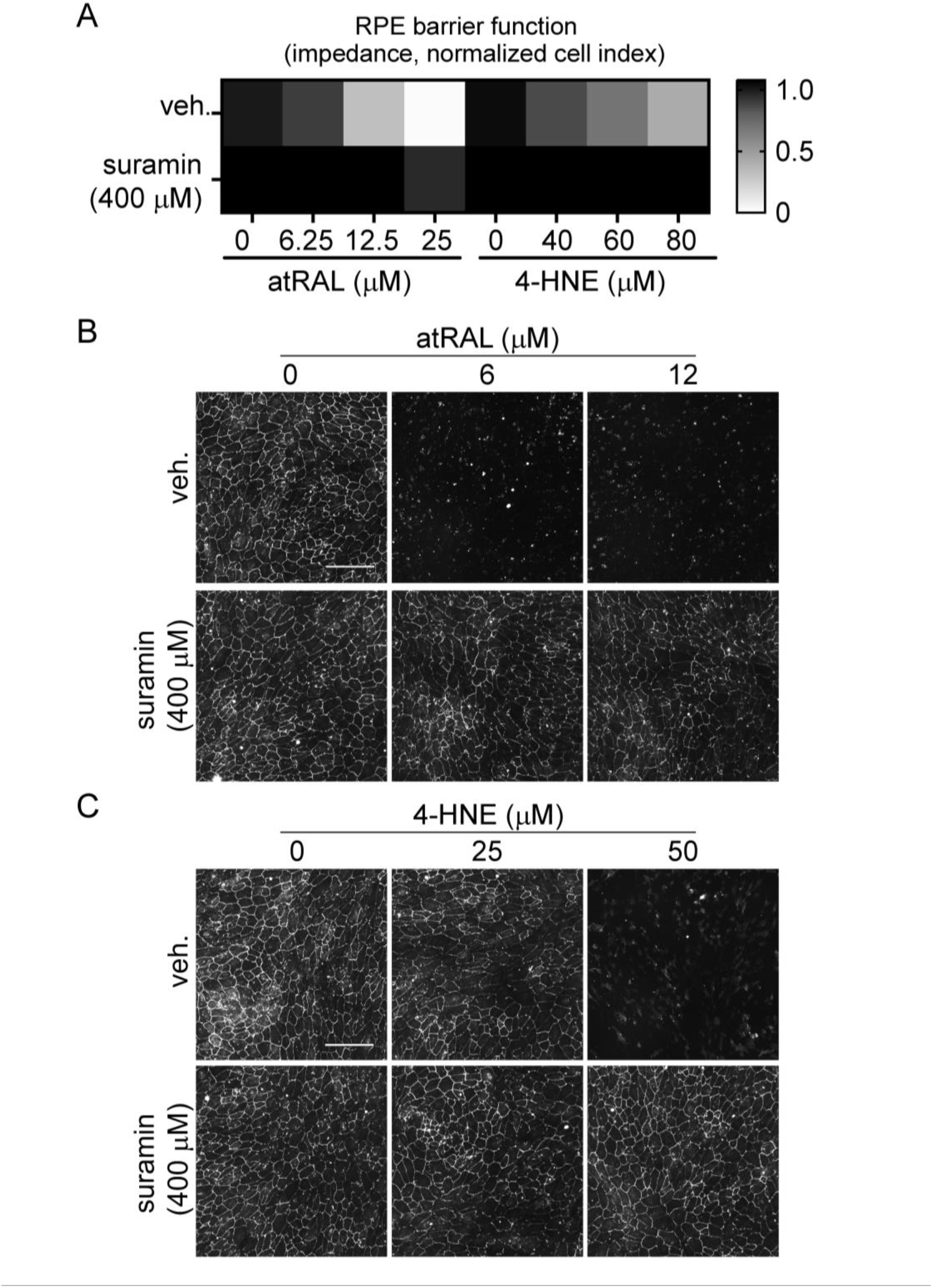
Suramin prevents toxin-induced ARPE-19 barrier function changes. (A) Suramin treatment of ARPE-19 exposed to atRAL or 4-HNE at different doses. RPE barrier function was evaluated via impendence measurements. Heat map indicates relative impedance to control. (B-C) Representative images of ZO-1 immunostaining after varying doses of atRAL (A) or 4-HNE (B) treatment with or without co-treatment with suramin shows the propertice effect of suramine on tocix-induced ZO-1 loss. Scale bar is 150 µm.

### Suramin shows protection from injury in zebrafish model

Purinergic signaling serves a key role in both PR and RPE health (19,36-39). In addition to its protective effect on RPE exposed to visual cycle metabolites, we sought to evaluate protective effects on PR. Although matured photoreceptors can be investigated *in vitro* using retinal organoid modelling, these lack a proper monolayer of RPE (40), thus we turned to a zebrafish model of RP to investigate the effect of purinergic receptor targeting on PR and RPE functions and survival. Transgenic zebrafish with GFP labeled OS were generated to express the RHO P23H mutation in rods, which has previously been reported to lead to a rapid loss of rod OS and subsequent cell death, as well as stress to the RPE (41-43). Mutant P23H zebrafish at 3 dpf were treated with either vehicle or 125 µM suramin, until 7 dpf, a timepoint at which rod photoreceptor degeneration is robust (38). Rapid and near complete loss of GFP+ PR were observed in the vehicle-treated group at 7 dpf (Figure 5A), which is consistent with previous report (43). However, suramin treatment led to a significant increase in GFP+ PR, compared to untreated control animals, indicating PR protection from degeneration (Figure 5A-B). Taken together, these results suggest that suramin inhibits intrinsic mechanisms of rod photoreceptor cell death caused by mutant RHO, possibly through its effect on purinergic signaling.

**Figure 5.**
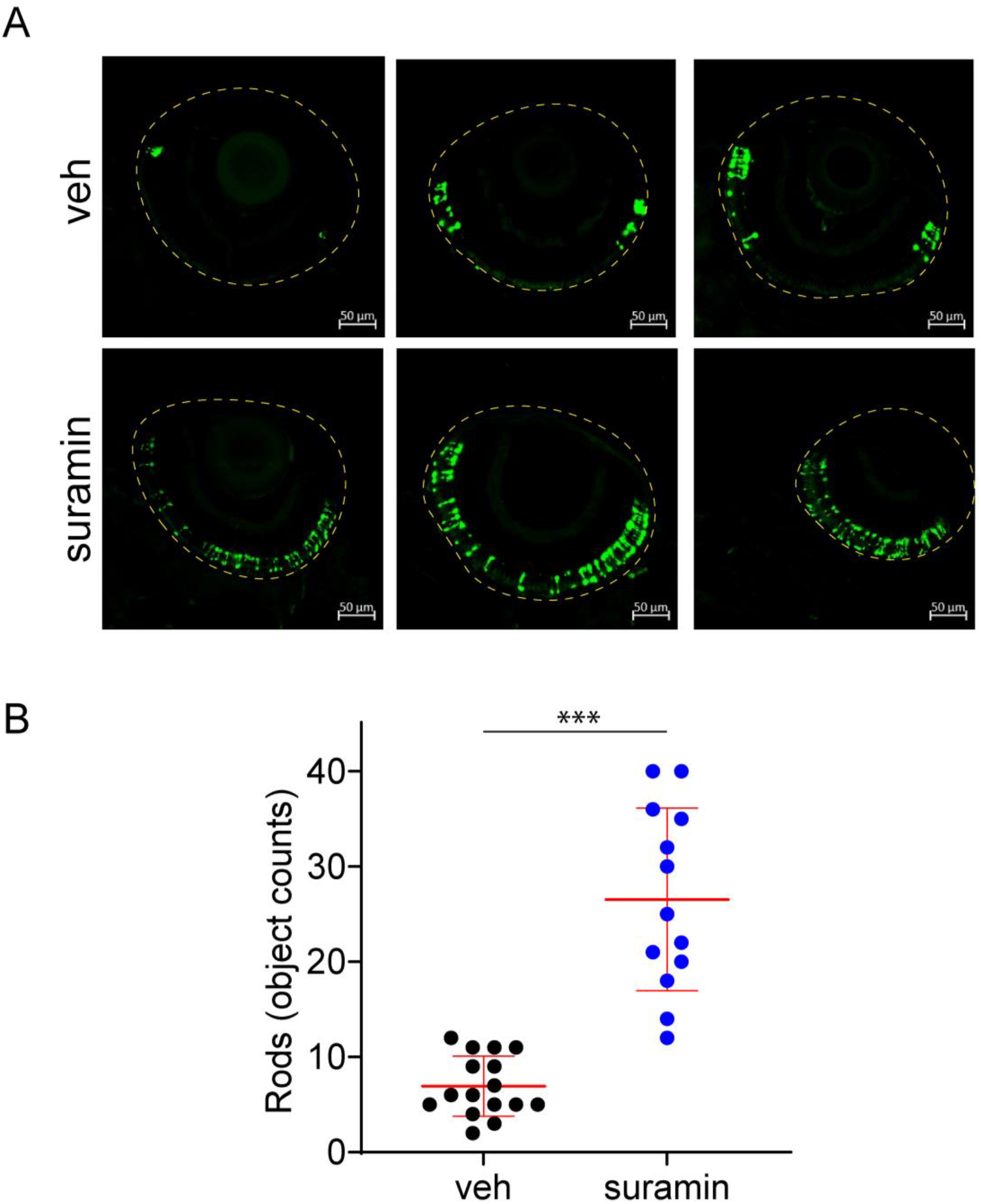
Suramin prevents photoreceptor loss in RHO-P23H zebrafish model. Retinas of RHO-P23H fish treated with vehicle or 125 µM suramin at 3 dpf until 7 dpf. Retinas were collected and stained by anti-GFP antibody. Remaining GFP+ rod were counted from least 4-5 animals per group, and the experiments were repeated 3 times. (A) Three representative images from each group of one experiment are presented. Scale bar is 50 µM. (B) Analyzed rod numbers were summarized from all animals. Data presented as mean±S.D., ***, *p*<0.001 compared to vehicle group.

## Discussion

ATP can be released from cells and lead to purinergic signaling via both autocrine and paracrine models. It was first described in neuronal cells and the central nervous system (44), and then widely observed in many other cell types, such as cancer cells and immune cells (45-49). Recently, evidence has suggested that ATP and purinergic signaling are regulating RPE functions and homeostasis, and are involved in retinal degenerative disorders (17,50). Increased ATP was observed in vitreous humor from AMD patients (16), and released ATP could cause RPE degeneration *in vitro* (19). Similar with this report (19), we found that extracellular ATP could induce RPE stress and functional decline, ultimately resulting in cell death. In addition, we also observed that extracellular ATP simultaneously sensitizes RPE to other stimuli-induced cell death (Figure 1). Inhibition of purinergic signaling using suramin and P2Y inhibitors, in addition to protecting RPE from stress induced cell death, also recovered metabolite-induced barrier function loss (Figure 3-4). These results highlight the therapeutic potential of purinergic receptor antagonism in preventing RPE degeneration and functional loss.

Suramin is a synthetic compound historically used to treat African sleeping sickness since 1922. The therapeutic potential for suramin to treat diseases ranging from autism to cancer has been well documented (51-55). As a pan-purinergic receptor antagonist, suramin can efficiently block different stimuli-induced RPE stress, functional loss and cell death, suggesting an important role of purinergic receptors in regulating RPE cell fitness. Many degenerative ocular diseases are commonly undiagnosed at early stages due to their slow progression and lack of clinical symptoms. The ideal treatment would be an intervention that can slow, halt, or even reverse the degenerative process in damaged cells. In order to evaluate if suramin could do this, we challenged cells with atRAL or 4-HNE with a time-course delay of treatment with suramin. Importantly, regardless of when suramin was introduced, protective effects were observed (Figure 2). The historically safe and efficacious use of suramin, combined with its flexible mode of intervention, makes further exploration of its therapeutic potential for ocular diseases attractive.

Pannexin-1 have been previously identified as a potential ATP release channel in different cell types (56,57). However, in our study, blockade of pannexin-1 using CBX only partially inhibited toxin-induced RPE cell death (Figure 3). This finding suggests that ATP release from RPE cells may not solely rely on the pannexin-1 channels. To further investigate the involvement of purinergic receptors in RPE cell death, we employed different compounds targeting various P2RY and P2RX receptors. Surprisingly, we discovered that blocking P2Y receptors using pan-P2Y antagonist could effectively prevent RPE cell death, whereas blocking P2X receptors using pan-P2X antagonist did not yield the same protective effects. It is worth noting that blocking each P2Y receptor individually was not sufficient to prevent cell death (Figure 3). These results suggest that pan-P2Y antagonism may hold therapeutic potential in the treatment of ocular diseases. Further understanding the complexity of ATP release from retinal cells would be an important step in developing such a strategy.

The data presented here supports a role of ATP and purinergic signaling in RPE degeneration *in vitro*. Protection of PR is equally as important for the maintenance of visual function. We explored the potential of suramin in preventing retinal degeneration *in vivo* using the RHO-P23H zebrafish model. This RP model carries a P23H mutation of RHO in rod photoreceptors, resulting in rapid and continuous rod degeneration (43). Rod loss was observable at 7 dpf, and this degeneration was ameliorated by suramin treatment. This *in vivo* study shows that blocking purinergic signaling could protect against genetic stress-induced retinal degeneration, such as that found in RP.

This study illustrates the effect of inhibiting purinergic receptors using suramin to prevent RPE and photoreceptor cell degeneration in multiple preclinical models, which give us strong confidence on our conclusion. Despite the caveats and challenges related to potential off-target effects of the compounds used and the redundancy of different purinergic receptors, our data strongly supports the involvement of ATP and purinergic signaling in RPE and retinal degeneration. This study highlights the potential of inhibiting purinergic receptors as a therapeutic strategy for retinal degenerative diseases.

## Ackowledgment

We thank Michael Capparelli (Global Discovery Chemistry, Biomedical Research, Novartis) for assistance with preparing 2-2Zn. The authors also thank to Tim Ramsey (Global Discovery Chemistry, Biomedical Research, Novartis) and Ganesh Prasanna (Ophthamology, Biomedical Research, Novartis) for his support and guidance on the project.

## Reference

1. Liao, Y., Zhang, H., He, D., Wang, Y., Cai, B., Chen, J., Ma, J., Liu, Z., and Wu, Y. (2019) Retinal Pigment Epithelium Cell Death Is Associated With NLRP3 Inflammasome Activation by All-trans Retinal. Invest Ophthalmol Vis Sci 60, 3034–3045

2. Somasundaran, S., Constable, I. J., Mellough, C. B., and Carvalho, L. S. (2020) Retinal pigment epithelium and age-related macular degeneration: A review of major disease mechanisms. Clin Exp Ophthalmol 48, 1043–1056

3. Yang, M., So, K. F., Lam, W. C., and Lo, A. C. Y. (2020) Novel Programmed Cell Death as Therapeutic Targets in Age-Related Macular Degeneration? Int J Mol Sci 21

4. Yang, S., Zhou, J., and Li, D. (2021) Functions and Diseases of the Retinal Pigment Epithelium. Front Pharmacol 12, 727870

5. Strauss, O. (2005) The retinal pigment epithelium in visual function. Physiol Rev 85, 845–881

6. Bonilha, V. L., Rayborn, M. E., Bhattacharya, S. K., Gu, X., Crabb, J. S., Crabb, J. W., and Hollyfield, J. G. (2006) The retinal pigment epithelium apical microvilli and retinal function. Adv Exp Med Biol 572, 519–524

7. Boulton, M., and Dayhaw-Barker, P. (2001) The role of the retinal pigment epithelium: topographical variation and ageing changes. Eye (Lond) 15, 384–389

8. Sparrow, J. R., Hicks, D., and Hamel, C. P. (2010) The retinal pigment epithelium in health and disease. Curr Mol Med 10, 802–823

9. Bowes Rickman, C., Farsiu, S., Toth, C. A., and Klingeborn, M. (2013) Dry age-related macular degeneration: mechanisms, therapeutic targets, and imaging. Invest Ophthalmol Vis Sci 54, ORSF68–80

10. Hanus, J., Anderson, C., and Wang, S. (2015) RPE necroptosis in response to oxidative stress and in AMD. Ageing Res Rev 24, 286–298

11. Tong, Y., and Wang, S. (2020) Not All Stressors Are Equal: Mechanism of Stressors on RPE Cell Degeneration. Front Cell Dev Biol 8, 591067

12. Hanus, J., Zhang, H., Wang, Z., Liu, Q., Zhou, Q., and Wang, S. (2013) Induction of necrotic cell death by oxidative stress in retinal pigment epithelial cells. Cell Death Dis 4, e965

13. Erb, L., Liao, Z., Seye, C. I., and Weisman, G. A. (2006) P2 receptors: intracellular signaling. Pflugers Arch 452, 552–562

14. Hasko, G., Linden, J., Cronstein, B., and Pacher, P. (2008) Adenosine receptors: therapeutic aspects for inflammatory and immune diseases. Nat Rev Drug Discov 7, 759–770

15. Fredholm, B. B., Ap, I. J., Jacobson, K. A., Linden, J., and Muller, C. E. (2011) International Union of Basic and Clinical Pharmacology. LXXXI. Nomenclature and classification of adenosine receptors--an update. Pharmacol Rev 63, 1–34

16. Notomi, S., Hisatomi, T., Murakami, Y., Terasaki, H., Sonoda, S., Asato, R., Takeda, A., Ikeda, Y., Enaida, H., Sakamoto, T., and Ishibashi, T. (2013) Dynamic increase in extracellular ATP accelerates photoreceptor cell apoptosis via ligation of P2RX7 in subretinal hemorrhage. PLoS One 8, e53338

17. Mitchell, C. H. (2001) Release of ATP by a human retinal pigment epithelial cell line: potential for autocrine stimulation through subretinal space. J Physiol 534, 193–202

18. Reigada, D., and Mitchell, C. H. (2005) Release of ATP from retinal pigment epithelial cells involves both CFTR and vesicular transport. Am J Physiol Cell Physiol 288, C132–140

19. Yang, D., Elner, S. G., Clark, A. J., Hughes, B. A., Petty, H. R., and Elner, V. M. (2011) Activation of P2X receptors induces apoptosis in human retinal pigment epithelium. Invest Ophthalmol Vis Sci 52, 1522–1530

20. Kurishita, Y., Kohira, T., Ojida, A., and Hamachi, I. (2012) Organelle-localizable fluorescent chemosensors for site-specific multicolor imaging of nucleoside polyphosphate dynamics in living cells. J Am Chem Soc 134, 18779–18789

21. Westerfield, M. (2000) The Zebrafish Book: A Guide for the Laboratory Use of Zebrafish (Danio Rerio), University of Oregon Press

22. Kwan, K. M., Fujimoto, E., Grabher, C., Mangum, B. D., Hardy, M. E., Campbell, D. S., Parant, J. M., Yost, H. J., Kanki, J. P., and Chien, C. B. (2007) The Tol2kit: a multisite gateway-based construction kit for Tol2 transposon transgenesis constructs. Dev Dyn 236, 3088–3099

23. Morris, A. C., Schroeter, E. H., Bilotta, J., Wong, R. O., and Fadool, J. M. (2005) Cone survival despite rod degeneration in XOPS-mCFP transgenic zebrafish. Invest Ophthalmol Vis Sci 46, 4762–4771

24. Kim, J. H., Lee, S. R., Li, L. H., Park, H. J., Park, J. H., Lee, K. Y., Kim, M. K., Shin, B. A., and Choi, S. Y. (2011) High cleavage efficiency of a 2A peptide derived from porcine teschovirus-1 in human cell lines, zebrafish and mice. PLoS One 6, e18556

25. Pearring, J. N., Lieu, E. C., Winter, J. R., Baker, S. A., and Arshavsky, V. Y. (2014) R9AP targeting to rod outer segments is independent of rhodopsin and is guided by the SNARE homology domain. Mol Biol Cell 25, 2644–2649

26. Meeker, N. D., Hutchinson, S. A., Ho, L., and Trede, N. S. (2007) Method for isolation of PCR-ready genomic DNA from zebrafish tissues. Biotechniques 43, 610, 612, 614

27. Uribe, R. A., and Gross, J. M. (2007) Immunohistochemistry on cryosections from embryonic and adult zebrafish eyes. CSH Protoc 2007, pdb prot4779

28. Maeda, A., Maeda, T., Golczak, M., and Palczewski, K. (2008) Retinopathy in mice induced by disrupted all-trans-retinal clearance. J Biol Chem 283, 26684–26693

29. Cia, D., Cubizolle, A., Crauste, C., Jacquemot, N., Guillou, L., Vigor, C., Angebault, C., Hamel, C. P., Vercauteren, J., and Brabet, P. (2016) Phloroglucinol protects retinal pigment epithelium and photoreceptor against all-trans-retinal-induced toxicity and inhibits A2E formation. J Cell Mol Med 20, 1651–1663

30. Li, J., Cai, X., Xia, Q., Yao, K., Chen, J., Zhang, Y., Naranmandura, H., Liu, X., and Wu, Y. (2015) Involvement of endoplasmic reticulum stress in all-trans-retinal-induced retinal pigment epithelium degeneration. Toxicol Sci 143, 196–208

31. Maeda, A., Maeda, T., Golczak, M., Chou, S., Desai, A., Hoppel, C. L., Matsuyama, S., and Palczewski, K. (2009) Involvement of all-trans-retinal in acute light-induced retinopathy of mice. J Biol Chem 284, 15173–15183

32. Bearelly, S., Khanifar, A. A., Lederer, D. E., Lee, J. J., Ghodasra, J. H., Stinnett, S. S., and Cousins, S. W. (2011) Use of fundus autofluorescence images to predict geographic atrophy progression. Retina 31, 81–86

33. Holz, F. G., Bellman, C., Staudt, S., Schutt, F., and Volcker, H. E. (2001) Fundus autofluorescence and development of geographic atrophy in age-related macular degeneration. Invest Ophthalmol Vis Sci 42, 1051–1056

34. Cai, B., Liao, C., He, D., Chen, J., Han, J., Lu, J., Qin, K., Liang, W., Wu, X., Liu, Z., and Wu, Y. (2022) Gasdermin E mediates photoreceptor damage by all-trans-retinal in the mouse retina. J Biol Chem 298, 101553

35. Kaarniranta, K., Ryhanen, T., Karjalainen, H. M., Lammi, M. J., Suuronen, T., Huhtala, A., Kontkanen, M., Terasvirta, M., Uusitalo, H., and Salminen, A. (2005) Geldanamycin increases 4-hydroxynonenal (HNE)-induced cell death in human retinal pigment epithelial cells. Neurosci Lett 382, 185–190

36. Fries, J. E., Goczalik, I. M., Wheeler-Schilling, T. H., Kohler, K., Guenther, E., Wolf, S., Wiedemann, P., Bringmann, A., Reichenbach, A., Francke, M., and Pannicke, T. (2005) Identification of P2Y receptor subtypes in human muller glial cells by physiology, single cell RT-PCR, and immunohistochemistry. Invest Ophthalmol Vis Sci 46, 3000–3007

37. Tovell, V. E., and Sanderson, J. (2008) Distinct P2Y receptor subtypes regulate calcium signaling in human retinal pigment epithelial cells. Invest Ophthalmol Vis Sci 49, 350–357

38. Corso, L., Cavallero, A., Baroni, D., Garbati, P., Prestipino, G., Bisti, S., Nobile, M., and Picco, C. (2016) Saffron reduces ATP-induced retinal cytotoxicity by targeting P2X7 receptors. Purinergic Signal 12, 161–174

39. Sanderson, J., Dartt, D. A., Trinkaus-Randall, V., Pintor, J., Civan, M. M., Delamere, N. A., Fletcher, E. L., Salt, T. E., Grosche, A., and Mitchell, C. H. (2014) Purines in the eye: recent evidence for the physiological and pathological role of purines in the RPE, retinal neurons, astrocytes, Muller cells, lens, trabecular meshwork, cornea and lacrimal gland. Exp Eye Res 127, 270–279

40. Singh, R. K., and Nasonkin, I. O. (2020) Limitations and Promise of Retinal Tissue From Human Pluripotent Stem Cells for Developing Therapies of Blindness. Front Cell Neurosci 14, 179

41. Goto, Y., Peachey, N. S., Ripps, H., and Naash, M. I. (1995) Functional abnormalities in transgenic mice expressing a mutant rhodopsin gene. Invest Ophthalmol Vis Sci 36, 62–71

42. Baid, R., Scheinman, R. I., Shinohara, T., Singh, D. P., and Kompella, U. B. (2011) LEDGF(1-326) decreases P23H and wild type rhodopsin aggregates and P23H rhodopsin mediated cell damage in human retinal pigment epithelial cells. PLoS One 6, e24616

43. Santhanam, A., Shihabeddin, E., Atkinson, J. A., Nguyen, D., Lin, Y. P., and O’Brien, J. (2020) A Zebrafish Model of Retinitis Pigmentosa Shows Continuous Degeneration and Regeneration of Rod Photoreceptors. Cells 9

44. Burnstock, G. (2008) Purinergic signalling and disorders of the central nervous system. Nat Rev Drug Discov 7, 575–590

45. Ledderose, C., Hefti, M. M., Chen, Y., Bao, Y., Seier, T., Li, L., Woehrle, T., Zhang, J., and Junger, W. G. (2016) Adenosine arrests breast cancer cell motility by A3 receptor stimulation. Purinergic Signal 12, 673–685

46. Ledderose, C., Bao, Y., Lidicky, M., Zipperle, J., Li, L., Strasser, K., Shapiro, N. I., and Junger, W. G. (2014) Mitochondria are gate-keepers of T cell function by producing the ATP that drives purinergic signaling. J Biol Chem 289, 25936–25945

47. Lee, A. H., Ledderose, C., Li, X., Slubowski, C. J., Sueyoshi, K., Staudenmaier, L., Bao, Y., Zhang, J., and Junger, W. G. (2018) Adenosine Triphosphate Release is Required for Toll-Like Receptor-Induced Monocyte/Macrophage Activation, Inflammasome Signaling, Interleukin-1beta Production, and the Host Immune Response to Infection. Crit Care Med 46, e1183–e1189

48. Chen, Y., Corriden, R., Inoue, Y., Yip, L., Hashiguchi, N., Zinkernagel, A., Nizet, V., Insel, P. A., and Junger, W. G. (2006) ATP release guides neutrophil chemotaxis via P2Y2 and A3 receptors. Science 314, 1792–1795

49. Bao, Y., Ledderose, C., Graf, A. F., Brix, B., Birsak, T., Lee, A., Zhang, J., and Junger, W. G. (2015) mTOR and differential activation of mitochondria orchestrate neutrophil chemotaxis. J Cell Biol 210, 1153–1164

50. Mitchell, C. H., and Reigada, D. (2008) Purinergic signalling in the subretinal space: a role in the communication between the retina and the RPE. Purinergic Signal 4, 101–107

51. Hough, D., Mao, A. R., Aman, M., Lozano, R., Smith-Hicks, C., Martinez-Cerdeno, V., Derby, M., Rome, Z., Malan, N., and Findling, R. L. (2023) Randomized clinical trial of low dose suramin intravenous infusions for treatment of autism spectrum disorder. Ann Gen Psychiatry 22, 45

52. Naviaux, R. K., Curtis, B., Li, K., Naviaux, J. C., Bright, A. T., Reiner, G. E., Westerfield, M., Goh, S., Alaynick, W. A., Wang, L., Capparelli, E. V., Adams, C., Sun, J., Jain, S., He, F., Arellano, D. A., Mash, L. E., Chukoskie, L., Lincoln, A., and Townsend, J. (2017) Low-dose suramin in autism spectrum disorder: a small, phase I/II, randomized clinical trial. Ann Clin Transl Neurol 4, 491–505

53. Wiedemar, N., Hauser, D. A., and Maser, P. (2020) 100 Years of Suramin. Antimicrob Agents Chemother 64

54. Villalona-Calero, M. A., Wientjes, M. G., Otterson, G. A., Kanter, S., Young, D., Murgo, A. J., Fischer, B., DeHoff, C., Chen, D., Yeh, T. K., Song, S., Grever, M., and Au, J. L. (2003) Phase I study of low-dose suramin as a chemosensitizer in patients with advanced non-small cell lung cancer. Clin Cancer Res 9, 3303–3311

55. Mirza, M. R., Jakobsen, E., Pfeiffer, P., Lindebjerg-Clasen, B., Bergh, J., and Rose, C. (1997) Suramin in non-small cell lung cancer and advanced breast cancer. Two parallel phase II studies. Acta Oncol 36, 171–174

56. Chen, Y., Yao, Y., Sumi, Y., Li, A., To, U. K., Elkhal, A., Inoue, Y., Woehrle, T., Zhang, Q., Hauser, C., and Junger, W. G. (2010) Purinergic signaling: a fundamental mechanism in neutrophil activation. Sci Signal 3, ra45

57. Schenk, U., Westendorf, A. M., Radaelli, E., Casati, A., Ferro, M., Fumagalli, M., Verderio, C., Buer, J., Scanziani, E., and Grassi, F. (2008) Purinergic control of T cell activation by ATP released through pannexin-1 hemichannels. Sci Signal 1, ra6

