## Supplementary information for "Protective effect of P2Y receptors antagonism on stress-induced retinal degeneration"

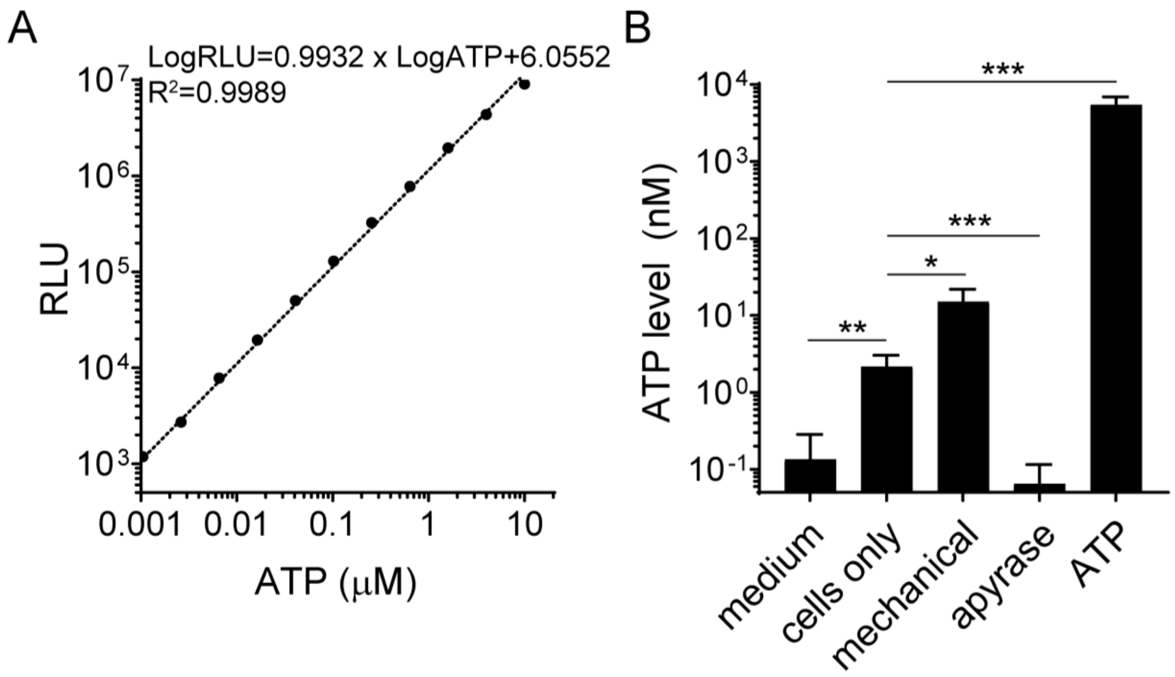


**Supplementary Figure 1. Extracellular ATP release was measured by luciferase-based ATP Determination Kit.** (A) Standard curve of ATP measurements. (B) Controls for measuring ATP in medium (no cells, blank control), cells only control, mechanical stimulation, apyrase (negative control) and ATP (10 µM, positive control) treated samples were measured. Relevant ATP amount was calculated based on the standard curve. Data presented as mean±S.D., * $p<0.05$, ** $p<0.01$ and *** $p<0.001$ compared to control.

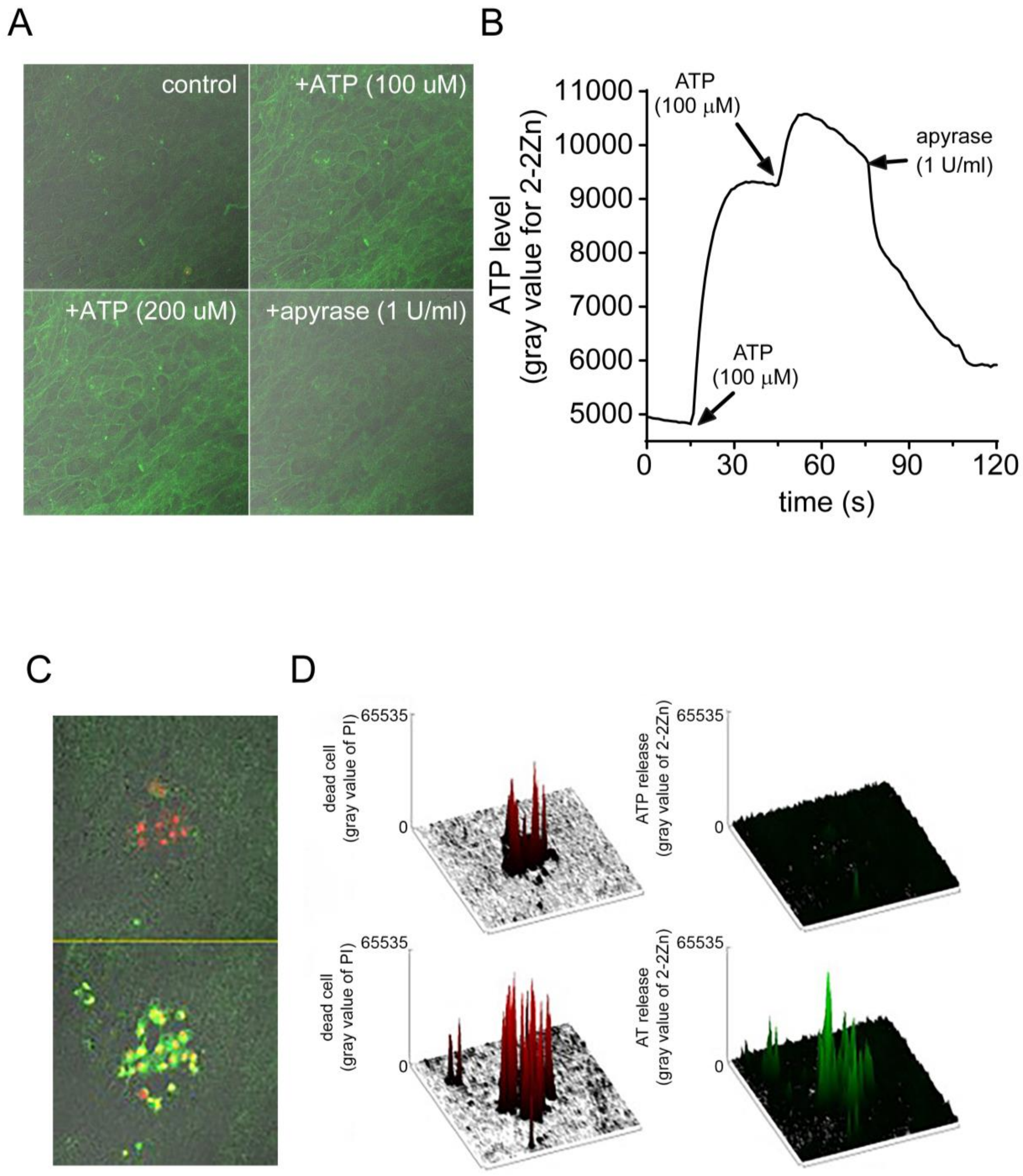


**Supplementary Figure 2. Visualization of ATP release from ARPE-19 cells under stress or death using membrane band ATP dye 2-2Zn.** Cells were treated with 1 μM 2-2Zn for 5 min. (A-B) ATP signal visualized using live cell imaging, also seen in Supplementary Movie 1. Basal levels of ATP release observed on cell membrane of ARPE-19, while enhanced with addition of ATP. Apyrase was used to chelate ATP and inhibit signal. Represented images were shown (A) and analyzed using Fiji software (B). (C-D) ATP release following laser-induced regional cell death, also seen in Supplementary Movie 2. Localized ARPE-19 cell death visualized by PI staining (red) immediately following laser exposure. Thirty min post-damage, a secondary cell death (red) and significant ATP release (green) were observed in ARPE-19.

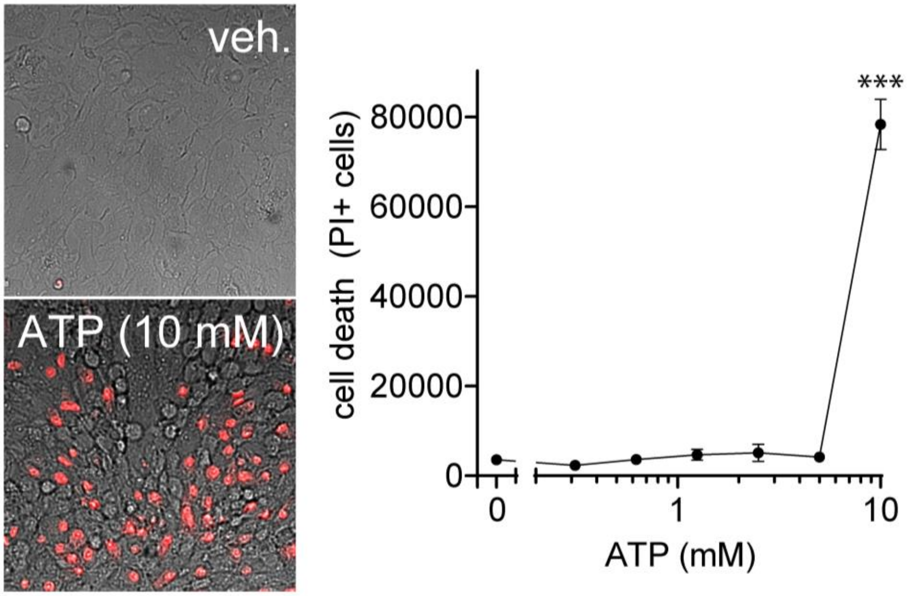


**Supplementary Figure 3. High dose ATP causes ARPE-19 cell death stained by PI.** ARPE-19 cells were treated with indicated doses of ATP, and cell death was evaluated by PI staining (red). Representative images for vehicle control and high dose of ATP (10 mM) are shown in the left panel. PI+ cells were quantified, and the dose response curve of ATP is shown on the right panel. Data presented as mean±S.D., ** $p$<0.01 compared to relevant control.
